# SimHS-AFMfit-MD: An Integrative Approach to Deciphering Alpha-Actinin Atomic Conformational Dynamics

**DOI:** 10.1101/2025.04.06.647477

**Authors:** Kien Xuan Ngo, Takashi Sumikama, Rémi Vuillemot, Han Gia Nguyen, Ngan Thi Phuong Le, Sergei Grudinin

**Author notes:** These authors contributed equally (Kien Xuan Ngo, Takashi Sumikama, Rémi Vuillemot, Sergei Grudinin). Corresponding author: Kien Xuan Ngo, Takashi Sumikama, Rémi Vuillemot, Sergei Grudinin, **Email:** (K.X.N.) (T.S.) (R.V.) (S.G.).

## Abstract

Many molecular systems, such as intrinsically disordered proteins and flexible multi-domain complexes, are highly dynamic and often inaccessible to conventional X-ray crystallography or cryo-EM due to their conformational heterogeneity and flexibility. As a result, resolving their atomic-level dynamics remains a significant challenge. In this study, we present SimHS-AFMfit-MD, an integrative framework that combines high-speed atomic force microscopy (HS-AFM), molecular dynamics (MD) simulations, and AFMfit-based structural modeling to reconstruct dynamic protein conformations at atomic resolution. Using alpha-actinin, an actin crosslinking protein, as a challenging test system, we show that AFMfit guided by nonlinear normal mode analysis (AFMfit-NMA) enables accurate structural fitting, while guiding AFMfit with MD trajectories (AFMfit-MD) further enhances the flexible fitting performance, achieving closer agreement with unbiased all-atom MD simulation results. This strategy allows us to convert thousands of three-dimensional HS-AFM images into atomic-scale conformational ensembles, revealing the twisting and bending transitions underlying Ca²⁺-bound and Ca²⁺-unbound alpha-actinin. Together, our results establish a hybrid computational–experimental approach that bridges the spatial and, to some extent, temporal resolution gaps between simulation and imaging, paving the way for real-time visualization of protein conformational dynamics at the atomic scale.

**Graphic Abstract:** 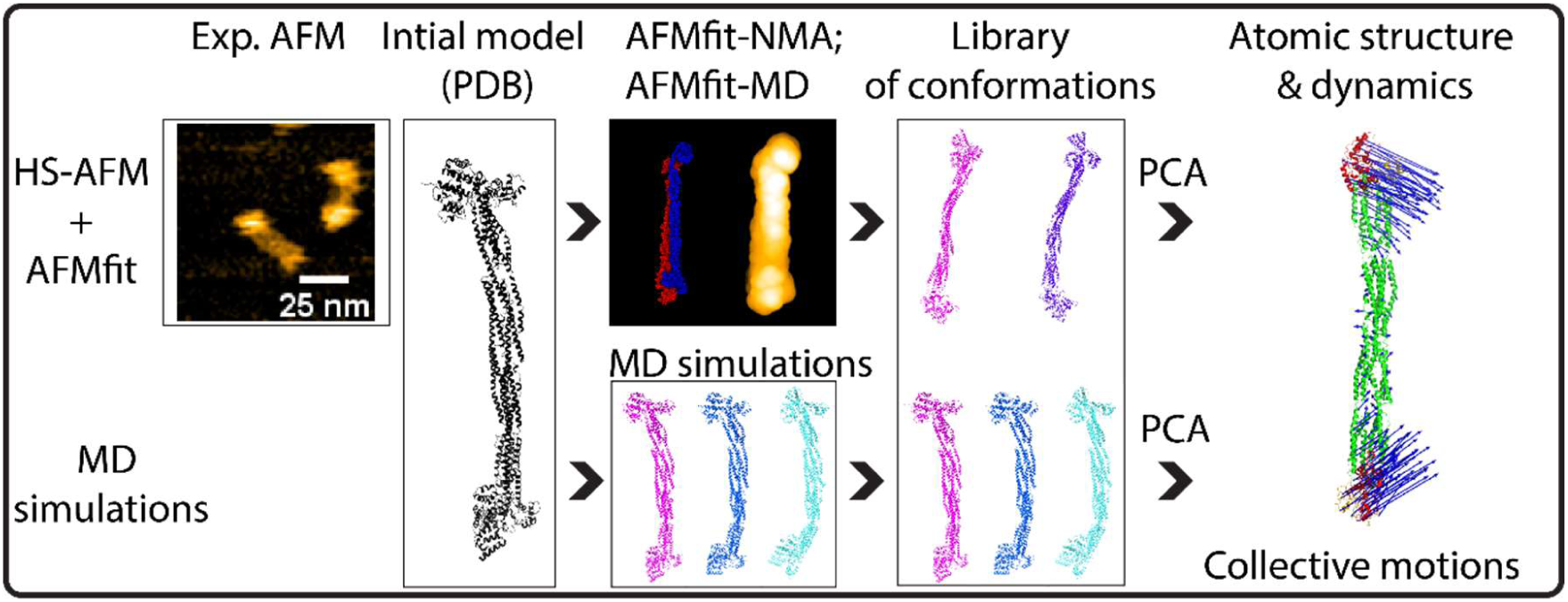

## Introduction

High-speed atomic force microscopy (HS-AFM) enables the imaging of biological structures and dynamics with nanometer-scale spatial resolution and millisecond temporal resolution. Generally, HS-AFM images capture three-dimensional (3D) surface data, incorporating two-dimensional (2D) lateral (x-y) and one-dimensional (1D) height (z) information encoded in pixel intensity^1^. This technique excels in imaging the conformational states, dynamics, and functional properties of both single and complex protein molecules, as well as intricate biomolecular conformations, offering capabilities beyond those of other imaging methods such as cryo-electron microscopy and X-ray crystallography, which primarily yield static structural data.

The primary challenge in studying protein dynamics at atomic resolutions in real time using atomic force microscopy (AFM) lies in bridging gaps at the temporal and spatial scales. While HS-AFM can capture the dynamics of the three-dimensional (3D) surface structures of proteins with optimal performance in just a few tens of milliseconds (∼20-100 ms/frame) and at nanometer scales (1 nm in the x‒y plane; ∼0.1 nm in the z‒direction)^1–11^, conventional molecular dynamics (MD) simulations typically explore atomic structures over much shorter time scales, from femtoseconds to microseconds^12–14^. Protein dynamics span a wide range of temporal scales (fs-µs), reflecting movements from atomic vibrations to molecular tumbling and larger collective motions that can be observed with nuclear magnetic resonance (NMR)^15,16^ and MD simulations^13,17–21^. However, HS-AFM is particularly valuable for investigating slower processes, such as protein folding and conformational changes induced by drugs or protein interactions, which occur over temporal scales ranging from milliseconds to seconds. Traditional MD simulations face challenges in modeling large and complex biological systems because of computational constraints. However, recent advancements have substantially extended their capabilities, allowing the simulation of larger biomolecular systems over longer timescales^22–25^. One of the alternatives to MD is the Normal Mode analysis (NMA), another classical technique, which is well-suited to study near-equilibrium protein dynamics. NMA represents protein motions in terms of independent components and can capture them on fs to µs temporal resolution. Importantly, modern NMA engines are ultra-fast, much faster than MD, and scale linearly with the system size^26^. HS-AFM complements these advancements by enabling high-speed imaging of complex protein surface structures and dynamics and provides detailed insights into their functional activities^1^. NMR can simultaneously resolve the atomic structures and interactional dynamics but remains limited by the protein size, even with recent advancements in in-cell NMR.

The very recent AFMfit-NMA method reconstructs 3D atomic-resolution structures by an iterative fitting process that progressively adapts an input model (e.g., that obtained from a protein data bank (PDB), an AlphaFold prediction, or a model from MD simulations) to match the 3D molecular surface images captured from multiple experimental AFM observations. AFMfit-NMA mutually explores the rigid and flexible degrees of freedom via the fast Fourier transform (FFT) and the nonlinear rigid block normal-mode analysis (NOLB-NMA) methods, respectively^26–28^. Overcoming the discrepancies in temporal and spatial resolution between the experimental AFM data and simulations would enable a thorough comparison of different experimental and computational techniques in dynamic atomic structural biology. To render such an analysis, we have chosen alpha-actinin, a very challenging molecular system that experiences large-scale motions both in temporal and spatial domains.

Alpha-actinin is a key protein in the organization of the actin cytoskeleton and is crucial for maintaining the mechanical properties of cells. In humans, alpha-actinin isoforms 2 and 3, which are believed to be Ca²⁺ insensitive, are integral to the function of striated muscle and are regulated by phosphoinositides^29–34^. In contrast, non-muscle alpha-actinin isoforms 1 and 4 are Ca^2+^-sensitive^35–37^. Previous studies have elucidated the atomic structure of these muscle and non-muscle alpha-actinin isoforms via techniques such as X-ray scattering^35–37^. However, these methods often fail to capture the dynamic conformational changes of these molecules. An understanding of these changes is crucial for elucidating the role of alpha-actinin in actin filament crosslinking and the associated structural changes under unipolar and bipolar crosslinking states.

In this study, we aimed to explore the atomic conformational dynamics of alpha-actinin through a multidimensional approach. To do so, we developed a novel workflow, SimHS-AFMfit-MD, that transforms the three-dimensional (3D) surface AFM images into 3D atomic conformations by integrating HS-AFM with the normal mode-based flexible fitting algorithm AFMfit-NMA^38^ and MD simulations^13,17–21^. This strategy bridges the spatial and, to some extent, temporal gaps between the experimental AFM data and MD/NMA simulations. Our pipeline enables the interpretation of the HS-AFM data with the 3D atomic models and overcomes the limitations of direct visualization techniques. MD simulations, together with the NMA, have extended our understanding of protein dynamics by capturing atomic structural transitions on the microsecond timescale, providing complementary insights into molecular flexibility and stability^39^.

## Results

### Overview of the methodology for integrating HS-AFM with AFMfit and MD simulations

The focus of this study was to develop an innovative workflow to analyze and understand how proteins change their atomic structure and how these conformational changes are related to their functions. Equally crucial, however, is understanding the protein dynamics, such as folding and interaction processes, since these processes directly influence the biological roles of proteins. HS-AFM can provide insights into the dynamic structural changes in proteins during their functional processes in solution^1,40,41^. While having high temporal resolution, HS-AFM lacks the atomic spatial resolution of other experimental techniques, such as X-ray crystallography or cryo-electron microscopy (cryo-EM). Therefore, one of the principal challenges with AFM lies in interpreting its low-resolution, 3D surface images into detailed 3D atomic conformations.

The innovative aspect of our approach lies in the combination of several techniques: HS-AFM, AFMfit (a method for fitting AFM images), nonlinear NMA, and MD simulations to establish the SimHS-AFMfit-MD pipeline (**Fig. 1**). This workflow consists of four key steps: (i) acquisition of experimental AFM images and protein models (PDB files); (ii-iii) AFM fitting (i.e. AFMfit-NMA, AFMfit-MD) and MD simulations to generate libraries of atomic conformations (PDB); and (iv) PCA analysis to classify the dynamic atomic conformations based on their principal components (PCs) (**Fig. 1C**). By integrating these techniques, we can obtain a more comprehensive view of the 3D atomic structure of proteins, along with their dynamics and conformational changes during functional activities.

**Fig. 1.**
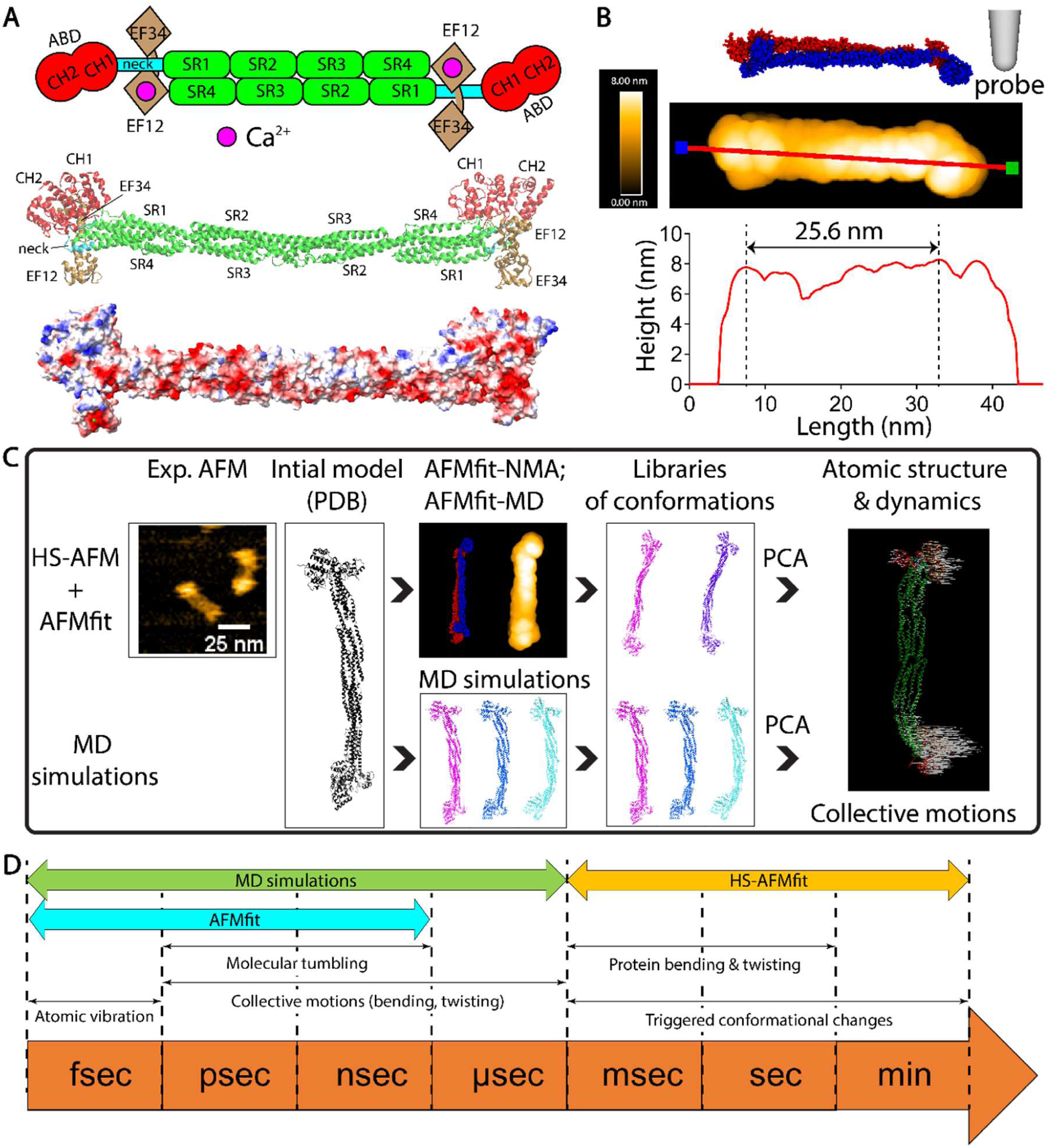
Novel workflow for analyzing the dynamics and atomic structures of alpha-actinin through the integration of HS-AFM with the AFMfit-NMA, AFMfit-MD, and MD simulations. (**A**) Domain composition and structure of chicken gizzard smooth muscle alpha-actinin (PDB: 1SJJ) depicted in a ribbon model and an electrostatic surface potential model. In the latter, red and blue represent negative and positive charges, respectively. Alpha-actinin is an antiparallel homodimer with a molecular weight exceeding 200 kDa. It comprises an N-terminal actin binding domain (ABD), in which the two calponin homology 1 and 2 (CH1, CH2) domains are in extensive contact, a central region containing four spectrin-like repeat (SR1-SR4) or rod domains, and a C-terminal calmodulin-like domain (CaMD) that includes two pairs of EF-hand motifs (EF12 and EF34). The EF12 domain is responsible for the Ca^2+^ binding. The two ABDs undergo a 90-degree rotation, enabling alpha-actinin to crosslink the actin filaments in nearly any orientation. (**B**) Pseudo-AFM images and topographic cross-sectional profile of alpha-actinin generated with the AFM viewer^75^. The pseudo-AFM images were obtained with the following scan parameters: step size, 0.1 nm; cone angle, 5°; and tip radius, 2.0 nm. The cross-sectional profile in the bottom panel corresponds to the red line in the middle panel. (**C**) Workflow for developing libraries of conformations, storing them as PDB files, and analyzing the dynamics and atomic structures of alpha-actinin through the integration of HS-AFM with the AFMfit-NMA or AFMfit-MD, and molecular dynamics (MD) simulations. This process is followed by principal component analysis (PCA) to identify and classify dynamic atomic structures into distinct modes of collective motion (indicated by white arrows). (**D**) Protein atomic conformational dynamics over timescales resolved by MD simulations and HS-AFM. Typically, protein conformational dynamics were resolved over nanosecond–second timescales using AFMfit, all-atom MD simulations, and HS-AFM.

We used our methodology to analyze the conformational changes in the Ca^2+^-bound and Ca^2+^-unbound alpha-actinin over milliseconds to seconds and to determine the atomic conformational dynamics of this protein. Alpha-actinin is an antiparallel homodimer with a molecular weight exceeding 200 kDa. The structure of chicken gizzard smooth muscle alpha-actinin (PDB: 1SJJ)^34^ consists of the following domains: an N-terminal actin binding domain (ABD), in which the two calponin homology 1 and 2 (CH1, CH2) domains are in extensive contact; a central region containing four spectrin-like repeat (SR1-SR4) or rod domains; and a C-terminal calmodulin-like domain (CaMD) that includes two pairs of EF-hand motifs (EF12 and EF34), and the EF12 domain is responsible for Ca^2+^ binding. The two ABDs undergo a 90-degree rotation, enabling alpha-actinin to crosslink the actin filaments in nearly any orientation^34^ (**Fig. 1A, B**).

We applied SimHS-AFMfit-MD to generate libraries of atomic models in different conformations for both Ca²⁺-bound and Ca²⁺-unbound alpha-actinin using a workflow shown in **Fig. 1C**. This provided insights into their atomistic structures and collective motions over time. First, we gently immobilized alpha-actinin molecules on mica and HS-AFM collected data with a typical imaging rate of 100 ms per frame (**Supplementary Video 1**). We then processed the HS-AFM with AFMfit-NMA or AFMfit-MD, starting with a reference atomic model derived from the MD simulations (**Supplementary Videos 2-3**). Specifically, AFMfit-NMA fits this reference model to the experimental AFM data by generating pseudo-AFM images derived from this reference model over two fitting steps. The first step was rigid fitting, where the reference model was rigidly aligned with the AFM experimental data to obtain an optimal pose using an exhaustive search algorithm based on the FFT. In the second flexible fitting step, the aligned models were flexibly deformed using the NOLB technique^26^ to obtain the optimal elastic alignment of the atomic models to multiple AFM observations. By applying this iterative process to our alpha-actinin AFM data, we collected the corresponding conformational libraries for the protein (see the Supplementary Methods). Simultaneously, the same initial protein model (PDB: 1SJJ)^34^ was used as the starting point for the MD simulations (see the Supplementary Methods). The simulations were conducted over a timescale of 1 microsecond, resulting in a set of conformation libraries for alpha-actinin. We then analyzed the atomic conformations obtained from AFMfit-NMA, AFMfit-MD, and MD simulations using principal component analysis (PCA). This analysis allowed us to interpret the conformational ensembles as representing the dynamic behavior of the atomistic structure of the proteins.

### Ca^2+^ binding affects smooth muscle alpha-actinin flexibility

First, we employed state-of-the-art HS-AFM to obtain high spatiotemporal resolution images of alpha-actinin. Three-dimensional molecular surface images of the individual Ca^2+^-bound and Ca^2+^-unbound alpha-actinin on mica were continuously recorded under near-physiological conditions at an imaging rate of 100 ms per frame (**Fig. 2, Supplementary Table 1, Supplementary Video 1**). The conformational dynamics of thousands of alpha-actinin molecules, including bending and twisting motions, were successfully captured and visualized in our AFM images (red and yellow arrowheads, **Fig. 2A**). To distinguish the conformational dynamics between these two states, we semi-automatically tracked the positions (x, y, and z coordinates) of the two ABD spheres on opposite sides of multiple alpha-actinin molecules over sequential frames and calculated the distance between their peak heights. HS-AFM analysis demonstrated that while the ensemble lengths of alpha-actinin were similar in the presence and absence of Ca²⁺, there was a significant difference in the ensemble heights between the two states. From the HS-AFM dataset, the mean ± SD heights and lengths were 5.7 ± 0.9 nm (N = 3040) and 25.1 ± 2.3 nm (N = 1941) for Ca^2+^-bound alpha-actinin and 4.8 ± 0.7 nm (N = 6080) and 25.0 ± 3.2 nm (N = 3885) for Ca^2+^-unbound alpha-actinin. The results revealed statistically significant differences (*p* < 0.05) in the height distributions, but no significant difference in length between the Ca^2+^-bound and Ca^2+^-unbound alpha-actinin (**Fig. 2B**).

**Fig. 2.**
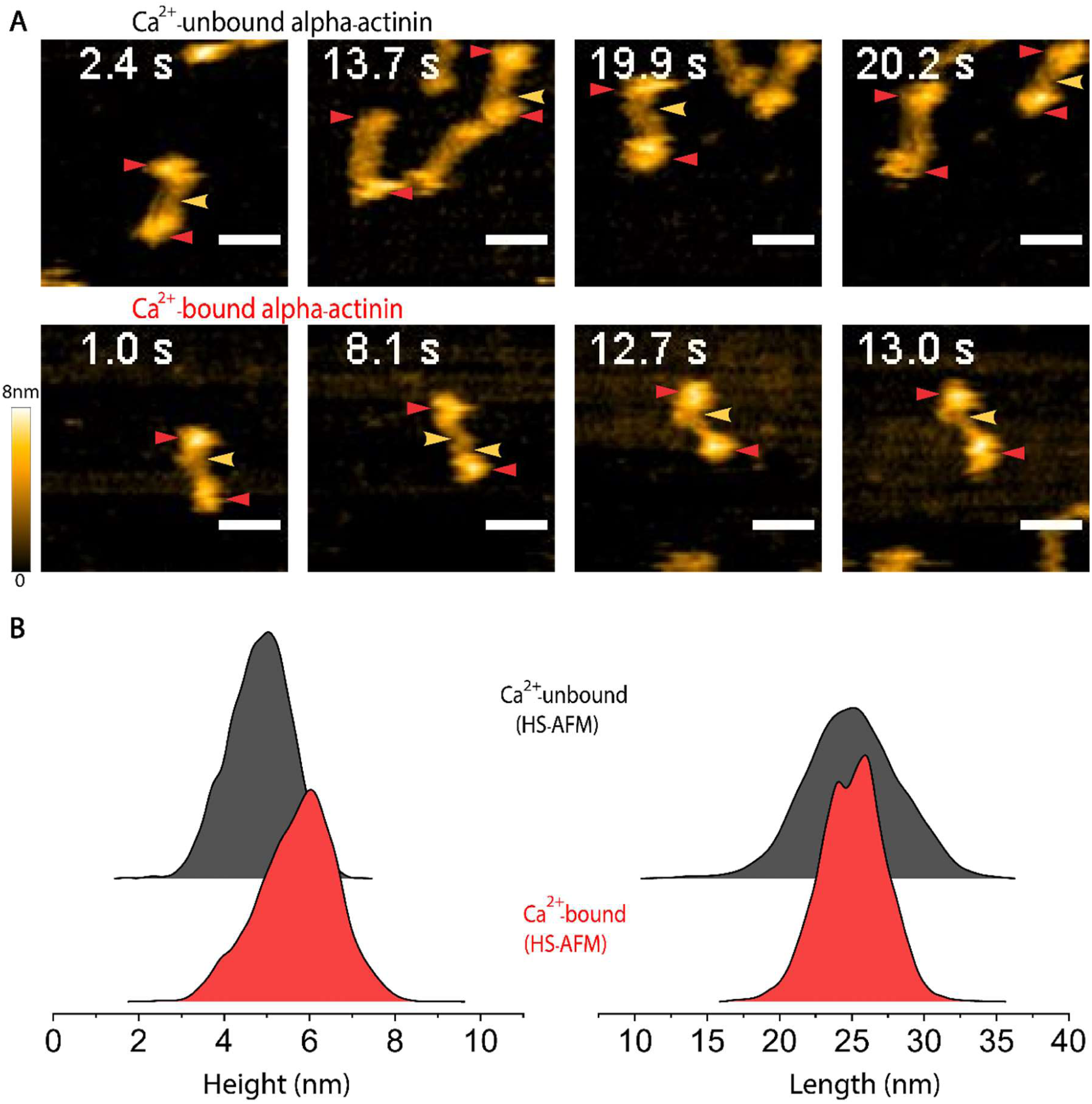
HS-AFM visualizes the 3D surface structure and millisecond conformational dynamics of alpha-actinin. (**A**) HS-AFM still images showing the various physical motions of alpha-actinin, such as twisting and bending, with and without Ca^2+^ binding. Yellow and red arrowheads denote bending and twisting motions, respectively, in the rod and neck domains. Scale bars: 25 nm. Color scale: 0–8 nm. (**B**) Ridgeline plots displaying the distribution of the peak heights and lengths between the two ABDs for Ca^2+^-bound (5.7 ± 0.9 nm, 25.1 ± 2.3 nm) and Ca^2+^-unbound (4.8 ± 0.7 nm, 25.0 ± 3.2 nm) alpha-actinin obtained with HS-AFM. A two-sample t-test (*p* ≤ 0.05) indicates that the difference in height distributions between the bound and unbound states is significant (*p* = 0), whereas the difference in length distributions is not significant (*p* = 0.78736). Related to **Supplementary Table 1**.

### Investigating alpha-actinin dynamics at the atomic level via SimHS-AFMfit-MD

Our HS-AFM–based analyses helped elucidate some of the characteristics of alpha-actinin, including its fluctuations in height and length over time. However, these conventional AFM processing techniques were unable to reveal critical insights into the conformational dynamics of alpha-actinin, such as its bending and twisting motions, as they only provided its surface structure (**Fig. 2A**). But an understanding of these motions is essential to grasp the mechanical influence of the actin-crosslinking function of alpha-actinin on the structure, dynamics, and function of actin filaments. Thus, we developed the SimHS-AFMfit-MD pipeline by integrating HS-AFM with AFMfit-NMA or AFMfit-MD, and MD simulations. This pipeline includes four steps (**Fig. 1C**) and allows us to further investigate the dynamics and atomic structure of alpha-actinin. We utilized AFMfit-NMA and AFMfit-MD to transform the 3D molecular surfaces of alpha-actinin, obtained from HS-AFM, into 3D atomic conformations (**Supplementary Fig. 1A, Supplementary Videos 2-3**). Through this method, we created libraries of atomic conformations for the Ca^2+^-bound and Ca^2+^-unbound alpha-actinin states (**Fig. 3A-C**). These conformations were then aligned to compare the dynamics of the two states based on the AFMfit-NMA, AFMfit-MD, and the MD simulations. The distribution of Cα positions was broader in the AFMfit-NMA analysis compared to the MD simulations (**Figs. 3A vs 3C**), yet the distributions were quite similar between AFMfit-MD and the MD simulations (**Figs. 3B vs 3C**). Notably, integrating MD trajectory information into AFMfit (AFMfit-MD) resulted in a conformational library that more closely aligned the one generated by MD simulations (**Figs. 3B vs 3C**). NMA-based structural fits, captured less extensive molecular change between Ca^2+^-bound and Ca^2+^-unbound states than the 1.0-microsecond unbiased MD simulations and AFMfit-MD. Additionally, substrate interactions that might have occurred during the AFM analysis, which were not present in the MD simulation results, could somehow influence the dynamics of alpha-actinin sitting on mica, even though immobilization conditions were optimized in our HS-AFM experiments.

**Fig. 3.**
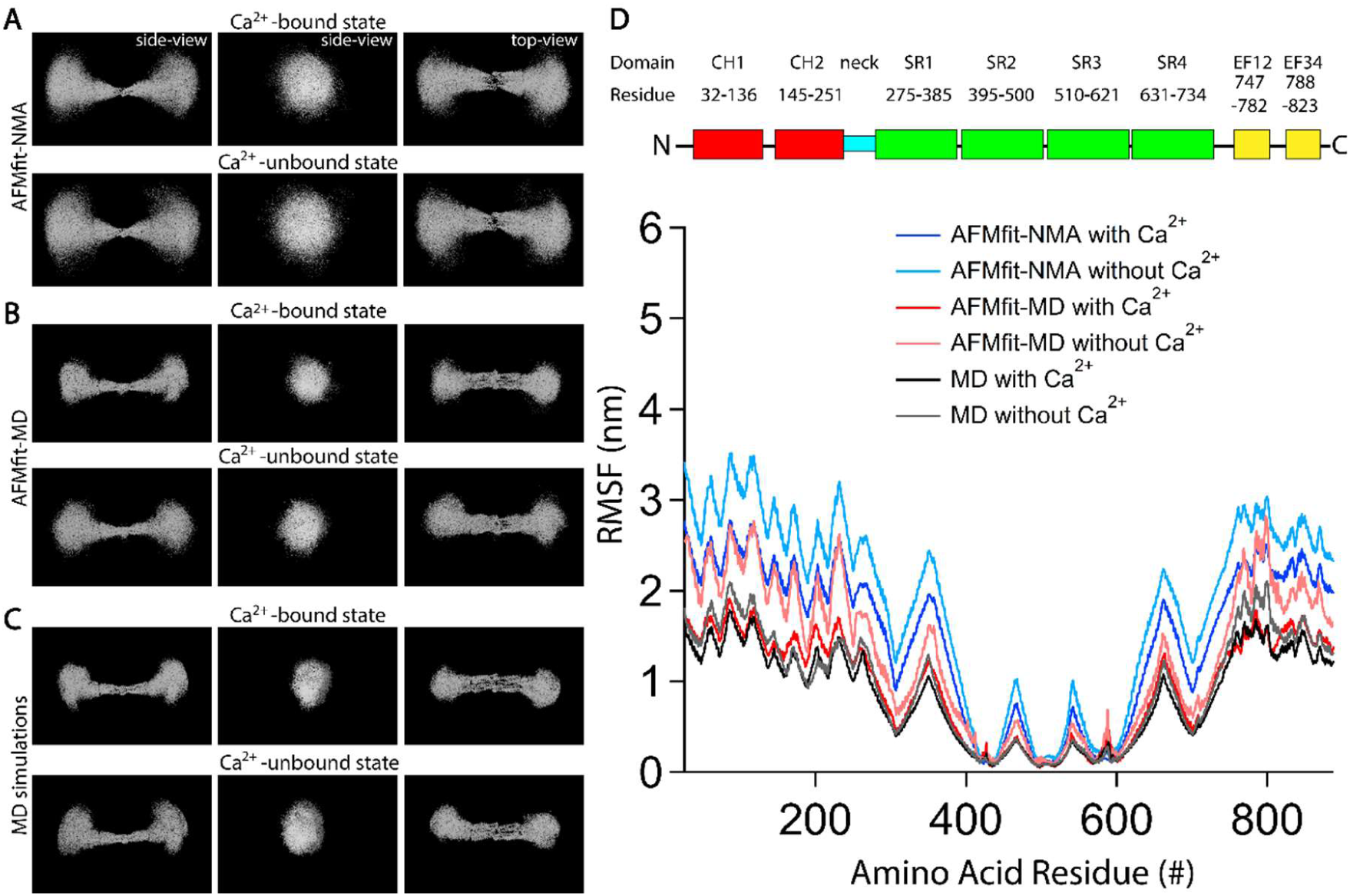
Analyses of the dynamics and atomic structure of the Ca^2+^-bound and unbound alpha-actinin using AFMfit-NMA, AFMfit-MD, and MD simulations. (**A, B, C**) Positions of the Cα atoms from the libraries of the Ca^2+^-bound and Ca^2+^-unbound alpha-actinin conformations obtained from the AFMfit-NMA, AFMfit-MD, and MD simulations, respectively. The conformations were aligned, assuming that the helical elements in the SR2 and SR3 domains (residues 485-525) exhibited the lowest root mean square fluctuation (RMSF). Top views and two different side views are presented to compare the molecular dynamics among the states and conformation libraries. (**D**) Root mean square fluctuation (RMSF) values plotted against the residue number. The illustration of the alpha-actinin domains, along with their corresponding amino acid residues, is based on PDB ID 1SJJ, using colors consistent with those in Fig. 1A. From the N-terminus to the C-terminus: CH1, CH2: calponin homology 1 and 2; SR1-SR4: spectrin-like repeat 1-4; and EF12, EF34: EF hand motifs 12, 34.

The root mean square fluctuation (RMSF), which is a commonly used metric for evaluating the flexibility and mobility of atoms or residues in proteins, was employed to quantify the deviation of each Cα atom from its average position using the conformational libraries generated with the AFMfit-NMA, AFMfit-MD, and MD simulations (**Fig. 3D**). The results revealed the dynamic motions of CH1-CH2 within the ABD, neck, and EF12-EF34 domains. However, the rod (SR1-SR4) domains exhibited relatively high rigidity.

Although there was a difference in the estimated fluctuation magnitudes of alpha-actinin between AFMfit-NMA and MD simulations, integrating MD trajectory guidance into AFMfit (i.e. AFMfit-MD) produced results that more closely aligned with those obtained from MD simulations. The SimHS-AFM-MD analysis consistently showed that Ca²⁺ binding slightly reduced the motion of the CH1-CH2 and EF12-EF34 domains near the neck in muscle alpha-actinin.

### Classifying the dynamic atomic conformations of alpha-actinin via PCA

PCA is a statistical method used to reduce the dimensionality of a dataset while retaining its most significant variations. It is highly effective in identifying and classifying the collective motions of individual protein domains using principal components (PCs).

First, we determined the ensemble of the atomic motions of the Ca²⁺-bound and Ca²^+^-unbound alpha-actinin by analyzing the libraries of the conformations obtained from the AFMfit-NMA, AFMfit-MD, and MD simulations and projected these motions onto a common PCA space (**Fig. 4A, Supplementary Videos 4-6**). The eigenvalues of principal components 1 (PC1), PC2, and PC3 were used to determine the percentage contribution of each motion. PC1 (27%) primarily captured the ensemble of the bending motions in the x-axis, PC2 (17%) represented the bending motions in the y-axis, and PC3 (10%) described the twisting motions.

**Fig. 4.**
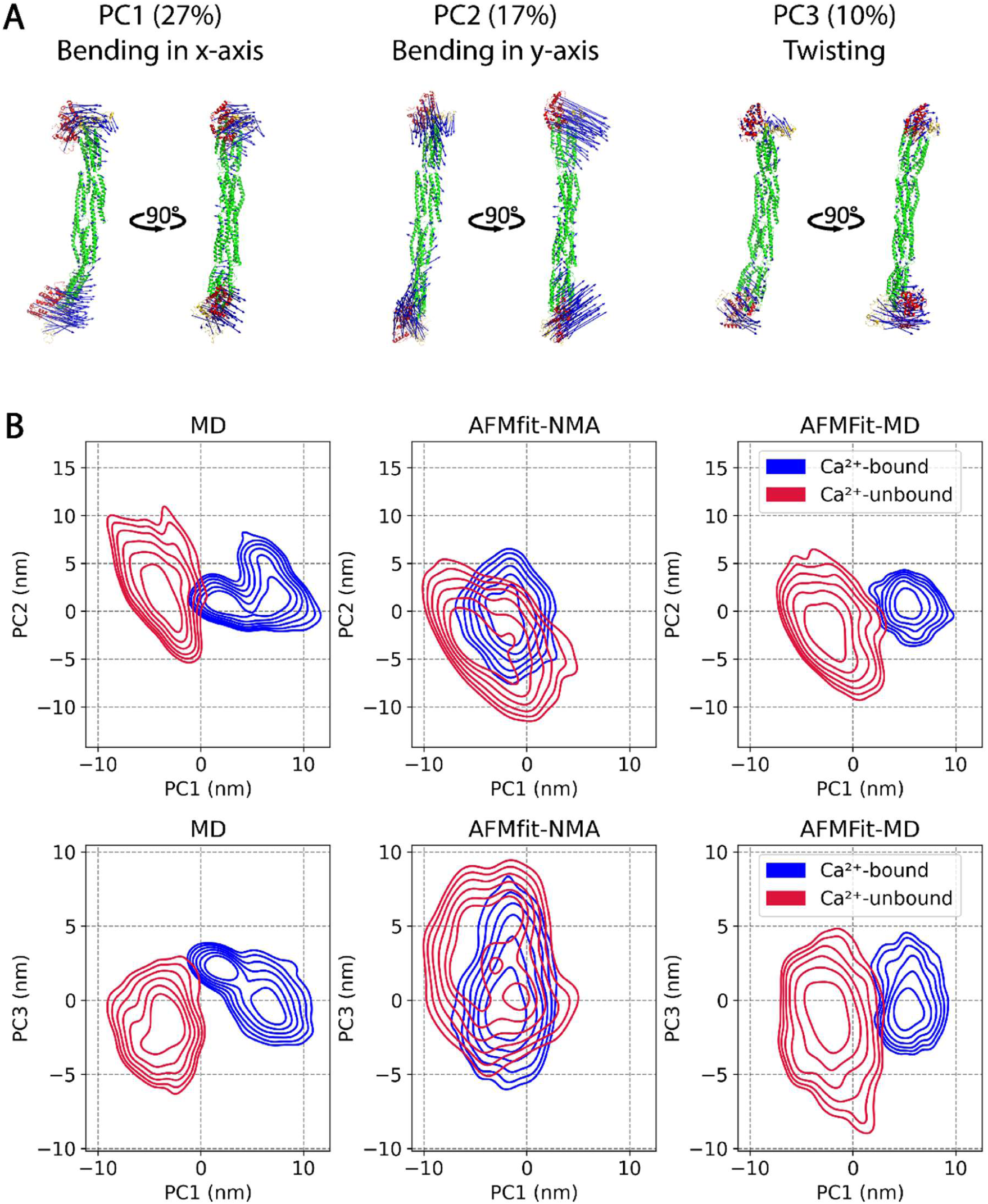
Analyses of the dynamics and atomic structure of the Ca^2+^-bound and Ca^2+^-unbound alpha-actinin using HS-AFM combined with the AFMfit-NMA or AFMfit-MD, and MD simulations. (A) Ensemble motions of the atomic conformations of alpha-actinin projected onto a common PCA space according to the conformational libraries of Ca^2+^-bound and Ca^2+^-unbound alpha-actinin obtained from the AFMfit-NMA, AFMfit-MD, and MD simulations. The eigenvalues of the principal components (PC1, PC2, PC3) were used to calculate the contributions (in percentages) of each motion to the total motions of alpha-actinin in the two states. The alpha-actinin structures are illustrated in the tube conformations, with the ABD, neck, rod, and EF12-EF34 domains represented in red, cyan, green, and yellow, respectively; the eigenvector components indicating the direction of motion are shown with blue arrows. (B) Distributions of PC1, PC2, and PC3 values. Data obtained from conformational libraries generated by AFMfit-NMA, AFMfit-MD, and MD simulations are shown in blue for the Ca²⁺-bound and in red for the Ca²⁺-unbound states of alpha-actinin.

To compare the data obtained from the AFMfit-NMA, AFMfit-MD, and MD simulations, we projected the conformational libraries of the Ca²⁺-bound and Ca²⁺-unbound alpha-actinin generated from the AFMfit-NMA, AFMfit-MD, and MD simulations onto a common PCA space and plotted the distributions of PC1 versus PC2 and PC1 versus PC3 (**Fig. 4B**). For both the Ca²⁺-bound and Ca²⁺-unbound states, PC1 derived from the AFMfit-NMA conformational libraries exhibited a broader distribution of ensemble bending motions along the x-axis compared with those from the AFMfit-MD and MD simulation libraries, which showed similarly narrower distributions.

Despite the differences in the fluctuation magnitudes exhibited by the AFMfit-NMA results and MD simulations, the integration of MD trajectory guidance into AFMfit produced a similar distribution of fluctuation (**Fig. 4B**), the PC2 value consistently captured the ensemble bending motions in the y-axis of both the Ca^2+^-bound and the Ca^2+^-unbound states. These results indicated the agreement between the three methods. We observed a similar trend for PC3, which captured a comparable degree of twisting motions in the Ca^2+^-bound and Ca^2+^-unbound states between AFMfit-MD and MD simulation methods. However, PC3 exhibited distinct fluctuations when compared to results obtained from AFMfit-NMA. Thus, we can conclude that PCA successfully analyzed and categorized the physical motions in the Ca^2+^-bound and Ca^2+^-unbound alpha-actinin at atomistic scales using PC1, PC2, and PC3, derived from the conformations generated via AFMfit-NMA and AFMfit-MD, analysis and the MD simulations (**Fig. 4**).

Across all results, the eigenvector components or motions represented by the blue arrows in SR1 and SR4 were small in magnitude. However, those in SR2 and SR3 showed almost no motion (**Fig. 4A**). These results confirmed the rigidity of the rod domain. In contrast, prominent eigenvector components were observed in CH1–CH2 within the ABD, in EF12– EF34, and in the neck domain, indicating relatively high flexibility in these regions. In summary, our SimHS-AFMfit-MD approach demonstrates an outstanding ability to capture and categorize the atomic conformational dynamics of the Ca²⁺-bound and Ca²⁺-unbound alpha-actinin.

### Quantitative analysis of domain distances in alpha-actinin

To more accurately characterize the dynamic atomic conformational changes in Ca²⁺-bound and Ca²⁺-unbound alpha-actinin, we conducted a quantitative analysis of the atomic-level movements of individual domains using AFMfit-NMA, AFMfit-MD, and MD simulations. We then compared the results across methods by focusing on the distances between the CH1, CH2, EF12, and EF34 domains within the ABD and CaMD (**Fig. 5, Supplementary Table 2**). This analysis allowed us to successfully capture the fluctuations between domains and assess the similarities and differences between the Ca²⁺-bound and Ca²⁺-unbound states.

**Fig. 5.**
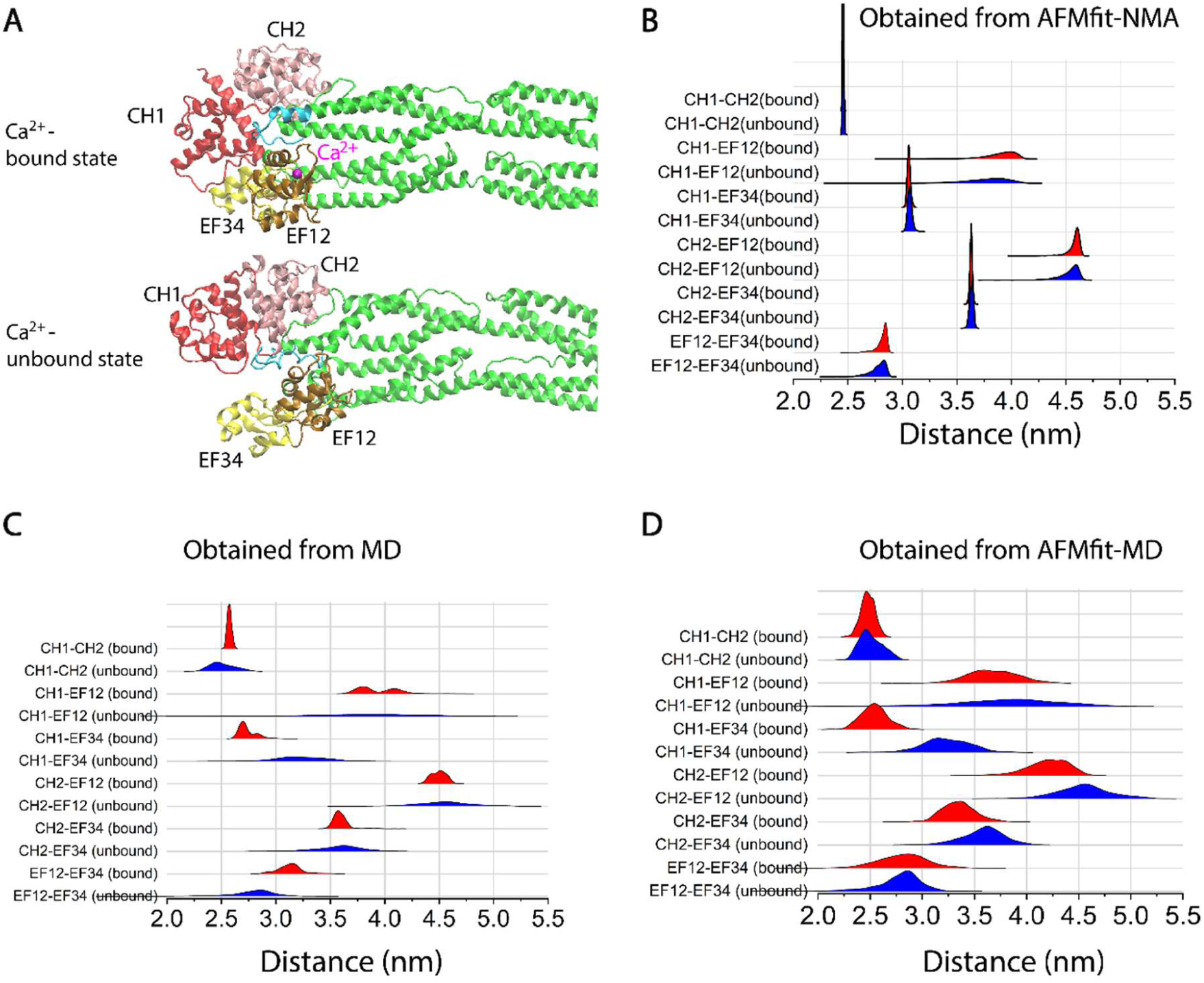
Analysis of domain distances in Ca²⁺-bound and Ca^2+^-unbound alpha-actinin based on conformations obtained from AFMfit-NMA, AFMfit-MD, and MD simulations. (A) Structural comparison of Ca²^+^-bound and Ca^2+^-unbound alpha-actinin, showing the positions of the CH1, CH2, EF12, and EF34 domains. (**B, C, D**) Distribution of distances between the CH1, CH2, EF12, and EF34 domains in Ca²^+^-bound (N = 1466) and Ca^2+^-unbound (N = 1140) alpha-actinin derived from AFMfit-NMA, AFMfit-MD (N = 1000), and MD (N = 1000) conformations, respectively. The domain distances were measured separately for chain 1 and chain 2 in individual conformations (stored as PDB files) for both Ca^2+^-bound and Ca^2+^-unbound states. The final data represent the averaged values of the two chains. Related to **Supplementary Table 2**.

The results from AFMfit-MD and MD simulations corroborated the molecular dynamics observed in the Ca²⁺-bound and Ca²⁺-unbound alpha-actinin structures obtained through AFMfit-NMA, revealing similar trends at the atomic level (**Fig. 5**). Notably, AFMfit-MD and MD simulations revealed greater magnitudes of domain fluctuations compared to AFMfit-NMA (**Figs. 5B vs 5CD**). The results from AFMfit-MD and MD simulations consistently showed that Ca²⁺ binding to the EF12 domain reduced fluctuations in alpha-actinin dynamics, whereas this effect captured via PC1 was not as clearly observed in the AFMfit-NMA results. In conclusion, our SimHS-AFMfit-MD pipeline successfully captured and characterized the atomic conformations and dynamic behaviors of both Ca²⁺-bound and Ca²⁺-unbound alpha-actinin across nanosecond to millisecond timescales, thereby bridging the spatial and, to some extent, temporal resolution gaps between experimental and computational approaches. This approach addresses the limitations of earlier static atomic structural studies that relied solely on X-ray scattering^33,35–37^ and cryo-EM^34^.

### Quantitative analysis of bending and twisting motions in alpha-actinin

To further capture the collective motions of the molecules and assess the physical impact of its actin crosslinking function, we analyzed the bending and twisting angles of the ABD relative to the rod domain in both the Ca^2+^-bound and Ca^2+^-unbound alpha-actinin using conformational libraries derived from AFMfit-NMA, AFMfit-MD, and MD simulations (**Fig. 6**). As described above, PCA of the conformations confirmed the rigidity of the rod domain (SR2-SR3). With reference to a vector defined along the Cα positions within the SR2-SR3 domain (**Fig. 6A**), the bending angles were measured relative to the center of mass of the Cα atoms in the CH1, CH2, EF12, and EF34 domains near the neck in chain 1 and chain 2. To measure the twisting angles, the vectors were defined by linking the centers of mass of the Cα atoms between the EF12-EF34 and CH1-CH2 domains, and the angles of these vectors were measured relative to the SR2-SR3-aligned reference vector (**Fig. 6B**). The distributions of bending and twisting angles in chain 1 and chain 2, with and without Ca²⁺ binding, were analyzed to more effectively capture the directions of the collective motions, whether in x-axis or y-axis direction. Quantitative measurements of the bending and twisting angles in chains 1 and 2 of the Ca²⁺-bound and Ca²⁺-unbound alpha-actinin based on the AFMfit-NMA and MD simulations (**Fig. 6, Supplementary Table 3**) confirmed the flexibility of the bending and twisting motions of alpha-actinin. Despite the previously identified differences between the methods in their ability to estimate the magnitudes of fluctuation, the AFMfit-NMA, AFMfit-MD, and the MD simulations showed that Ca²⁺ binding could restrict these motions to some extent. Our approach effectively refined and categorized the impact of Ca²⁺ on physical motions, such as the bending and twisting of smooth muscle alpha-actinin. When the ensemble of conformations obtained using our method was compared with those obtained from other techniques, such as cryo-EM^6^, which reported that non muscle alpha-actinin was not sensitive to Ca²⁺, our results not only aligned with these findings but also offered a more detailed understanding of its dynamic behavior than cryo-EM.

**Fig. 6.**
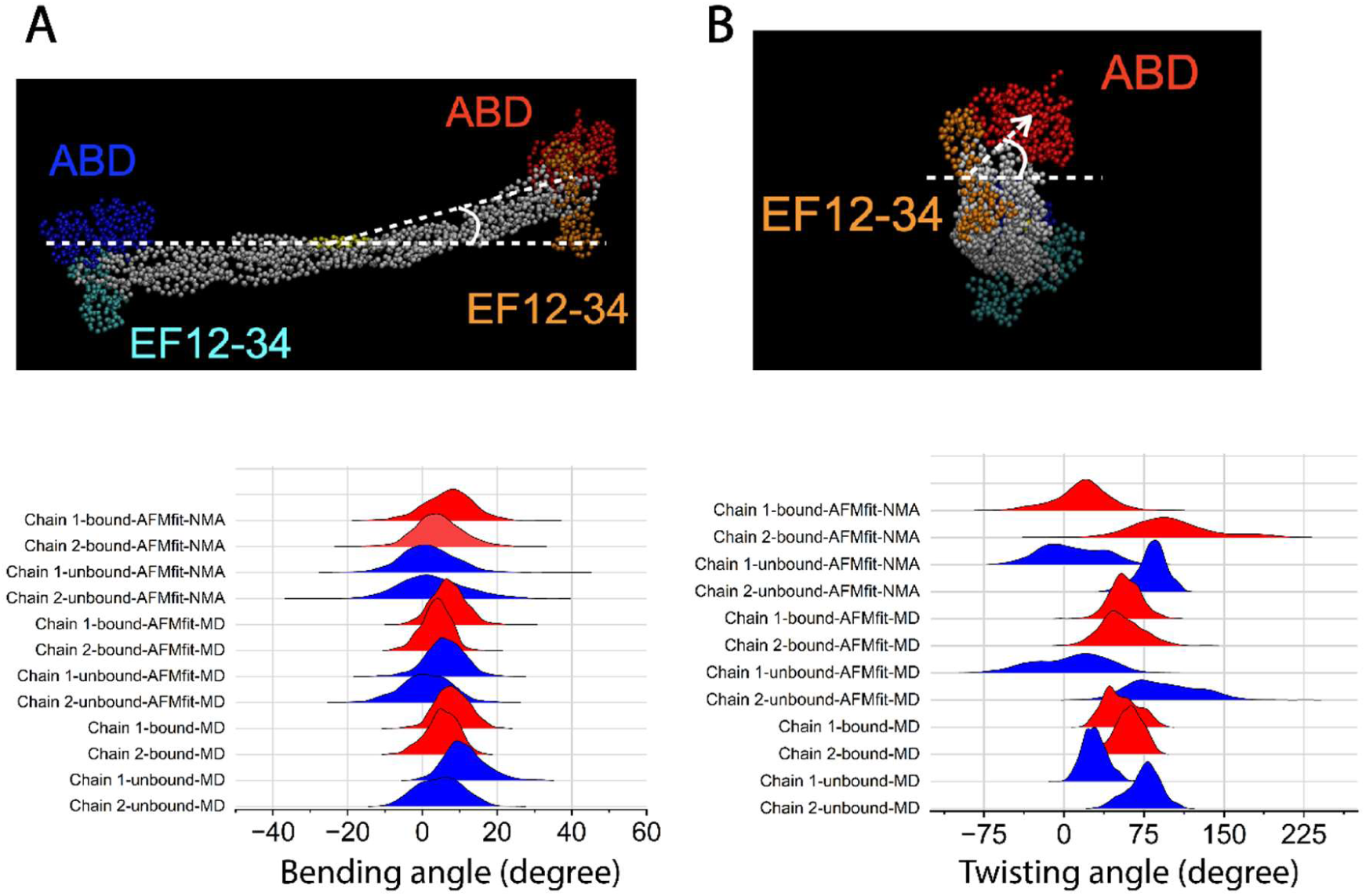
Analysis of the bending (A) and twisting (B) angles of the ABD relative to the rod domain of Ca^2+^-bound and Ca^2+^-unbound alpha-actinin using the conformations obtained from AFMfit-NMA, AFMfit-MD, and MD simulations. Related to **Supplementary Table 3**.

## Discussion

Time-resolved cryo-EM is emerging as a powerful tool for investigating protein dynamics and atomic conformations simultaneously^42–44^. Its potential can be further enhanced by integrating approaches such as cryo-EM flexible fitting^45^ or computational methods. Additionally, advanced NMR spectroscopy provides complementary insights and captures both structural and dynamic aspects of proteins during interactions^15,16,46,47^.

While NMR spectroscopy excels in studying protein structure, dynamics, and interactions, it has limitations. Traditional solution-state NMR is most effective for proteins under ∼30–50 kDa, although techniques such as transverse relaxation-optimized spectroscopy (TROSY) extend this range to ∼100 kDa. Larger proteins suffer from the signal overlap and line broadening because of the slow molecular tumbling and transverse relaxation. Compared with cryo-EM and X-ray crystallography, NMR also has lower sensitivity, and peak overlap complicates resonance assignments in complex systems^16,48–50^.

In dynamic studies, NMR provides insights into timescales from picoseconds to seconds but struggles with motions faster than 10 ps or slower than seconds to minutes^49,50^. Solid-state NMR remains challenging for membrane proteins because of the need for extensive isotopic labeling and specialized pulse sequences^50^. Additionally, weak or transient interactions can produce signals that are too faint for accurate characterization. Sample preparation and long data acquisition times (hours to days) further risk protein degradation or aggregation^48^.

While cryo-EM surpasses NMR in resolving large protein complexes (>100 kDa), it is less effective for studying dynamics^51^. However, these techniques can complement each other^52^. Other single-molecule techniques, such as FRET, can complement NMR for fast dynamics^53^, but no standalone method can perfectly capture the atomic structures, dynamics, and interactions simultaneously.

Despite the inherent limitations of AFM, such as its ability to capture only 3D surface images of soft and biological samples with a maximum spatial resolution of ∼1 nm in the x‒y plane and ∼0.1 nm in the z dimension, recent advancements in HS-AFM have significantly increased its temporal resolution^1^. Due to these improvements, imaging at timescales of a few tens of milliseconds or faster can be performed and enable a more precise characterization of protein dynamics. HS-AFM has the ability to image large complex protein systems without the need for chemical labeling, enabling the simultaneous capture of nanostructures, dynamics, and protein interactions or assemblies under near-physiological conditions; this capability surpasses those of NMR, x-ray crystallography, and cryo-EM. Additionally, by integrating HS-AFM with the AFMfit-NMA^38^ and AFMfit-MD and MD simulations through SimHS-AFMfit-MD, we generated comprehensive libraries of atomic conformations, facilitating the investigation of the physical motions and atomic structures of alpha-actinin through PCA and providing a robust framework for improving the interpretation of low-resolution experimental AFM data. For instance, the twisting and bending motions of alpha-actinin were cross-validated using AFMfit-based NMA and AFMfit guided by MD trajectories and unbiased all-atom MD simulations, confirming the robustness of the integrated SimHS-AFMfit-MD approach and offering deeper insights into the physical dynamics and actin crosslinking function of alpha-actinin. This capability potentially contributes to better understanding of previous static structural studies using X-ray diffraction^54,55^ and cryo-EM^56,57^. Ultimately, this integration addresses the spatial and temporal limitations of experimental AFM and MD simulation data and facilitates the study of the atomic conformational dynamics of complex protein assemblies. Apparently, the data obtained through SimHS-AFMfit-MD can be useful for developing force fields to simulate complex protein models, as demonstrated in previous studies^22–25^. Force fields are crucial for accurately validating the interatomic forces acting between the atoms composing the system since they account for the intermolecular or interatomic interactions and energy minimization^58,59^. More broadly, the conformational libraries could also serve as training data for the deep learning models^60–62^ to predict the pseudoatomic structures across different timescales (i.e., prediction of protein dynamics) and help bridge the gap between the experimental and computational methods. In this context, recent advancements in three-dimensional localization AFM (3D-LAFM) have successfully transformed the AFM data into 3D density data^63^, facilitating large-scale 3D atomic AFM data generation through the integration with the MD simulations.

## Conclusion

SimHS-AFMfit-MD provides an innovative approach for overcoming the spatial and temporal resolution limitations of imaging and simulation data. This method can be further developed, such as by its integration with deep learning, NMR, cryo-EM, cryo-ET, or in silico AFM^64^, to bridge the spatiotemporal resolution gaps between HS-AFM and the MD simulations and to capture and predict protein dynamics; this has enabled real-time studies of the atomic conformational dynamics of complex proteins and surpassing the capabilities of individual techniques. Our approach facilitates new possibilities for advancing the understanding of the atomic structural dynamics of complex protein assemblies that mimic cellular machines.

## Methods

### Proteins

Chicken gizzard smooth muscle alpha-actinin (isoform 2)^33^ was obtained from Cytoskeletons (Sigma, Japan).

### HS-AFM Analysis

We employed high-speed atomic force microscopy (HS-AFM), as previously described^7,8,40,65^, to capture the dynamic conformations of alpha-actinin in various states, providing high-speed imaging and three-dimensional (3D) surface structure of the protein at nanometer resolution. Chicken gizzard smooth muscle alpha-actinin, an isoform 2 (Sigma, Japan), was diluted to 30 nM and imaged in F-buffer containing 20 mM KCl, 10 mM PIPES-KOH (pH 6.8), 0.5 mM MgCl_2_, 0.25 mM EGTA, 0.25 mM DTT, and 0.5 mM ATP. To observe the dynamic conformational changes in alpha-actinin with CaCl_2_, the proteins were imaged in the same F-buffer with the addition of 0.1-0.2 mM CaCl_2_. Length and height of alpha-actinin were semi-automatical analyzed using Kodec software with the source code^7^.

### AFMfit-NMA Analysis

The AFMfit-NMA protocol was originally developed to fit 3D atomic structures into 3D surface AFM images^38^. In this study, AFMfit-NMA started the fitting procedure with the initial models used in MD simulations. These models were derived from PDB entry 1SJJ^34^ for both Ca²^+^-bound and Ca^2+^-unbound states, serving as a reference atomic model.

The AFMfit-NMA protocol began by cropping AFM images into sub-images, referred to as particles, that corresponded to the molecule of interest. This cropping operation was semi-automatic: an initial automated process identified potential particles using the particle-tracking toolkit TrackPy (Allan, D. B., Caswell, T., Keim, N. C., van der Wel, C. M., & Verweij, R. W. (2021). soft-matter/trackpy: Trackpy v0.5.0 (v0.5.0). Zenodo. https://doi.org/10.5281/zenodo.4682814), followed by a manual verification step to eliminate any false positives.

We processed the extracted particles using AFMfit-NMA, which estimated the optimal rigid and flexible degrees of freedom for the initial model. The alignment procedure maximized the similarity between the particles and the pseudo-AFM images calculated from the model. The calculation of these pseudo-AFM images had to be adjusted to match the experimental conditions. Specifically, we estimated the shape of the experimental probe tip to account accurately for the tip convolution effect. We conducted this estimation through an exhaustive search of a smoothness parameter across the entire set of particles.

AFMfit proceeded in two steps. In the first step, it performed a rigid-body pre-alignment of the model to the particles using a 5D exhaustive search accelerated by FFT. This process estimated the relative position (two translational degrees of freedom) and orientation (three rotational degrees of freedom) of each particle. The angular step for the rotational search was set to 10° and restricted to flat orientations of alpha-actinin to accelerate the computations. The translational step in the z-direction was set to 1 Å.

In the second step, AFMfit combined the continuous rigid and flexible alignments of the model to the particle using the non-linear normal mode analysis method NOLB^26^. NOLB calculated normal modes using the Rotation-Translation of Blocks (RTB) approximation on an Elastic Network Model (ENM). We set the interaction cutoff in the ENM to 8 Å. We restricted the possible motions of our model to the sixteen lowest frequency normal modes, among which ten modes were the flexible motions describing the most collective motions of alpha-actinin, including bending and twisting. The six normal modes left corresponded to rigid translations and rigid rotations in 3D. We used them to account for continuous rigid rearrangement that could be necessary when flexibly deformating the model. According to the AFMfit-NMA method, we iteratively calculated the optimal linear approximation of the normal mode displacement that fitted the particle. Then, we carried out a non-linear extrapolation of the motion following the NOLB methodology. We iterated over the linear approximation and non-linear extrapolation until convergence, which, in practice, was obtained in five iterations.

The flexible fitting procedure aimed to keep a balance between minimizing deformations from the initial model – thus preserving its structure and preventing unrealistic distortions – and maximizing similarity to the data. This balance was controlled by a hyperparameter λ. We estimated the optimal value of λ through an exhaustive search, conducting flexible fitting on a sample of particles. Our goal was to achieve a maximum average root-mean-square deviation (RMSD) of 10 Å from the initial model. We chose this threshold to reflect the typical range of motions observed in molecular dynamics (MD) simulations of alpha-actinin. Ultimately, we found that the optimal value for λ was 7.5 (**Supplementary Fig. 1**).

### AFMfit-MD

The original AFMfit method (we refer to it here as AFMFit-NMA) employed NMA and the elastic network model to estimate the low-dimensional subspace of protein motions. In the current AFMfit-MD algorithm, we utilize presampled MD conformations for estimating this subspace instead of the NMA. Concretely, we do the following. Firstly, we structurally align all the MD frames on the reference one by minimizing the corresponding RMSD using the QCP library^66^. We use the first MD frame as a reference. Then, we *linearly* project all the frames into the RTB six-dimensional SE(3) space, constructed at the reference conformation of the system. By default, we are using each residue as a rigid block. After, we build a 6*B* × 6*B* covariance matrix in this space, similarly to the RTB-projected Hessian matrix in the RTB NMA approaches^66^. Eigenvectors of this matrix correspond to the instantaneous linear and angular velocities of each rigid block, protein’s residue in our case. We shall mention that if the number of frames in a trajectory is much smaller than *6B*, it is computationally more advantageous to instead compute SVD vectors on the original matrix of RTB projected coordinates. Then, we proceed with the NOLB *nonlinear* extrapolation of the computed eigenmotions and the AFMfit fitting procedure.

### AFMfit-MD theory

Let us consider a rigid body parameterized with a 3-vector of its centre of mass *c⃗*, a 3-vector of its orientation 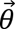, its total mass *m*, and its *inertia tensor I*, computed relatively to the center of mass. Assuming that it is composed of *k* individual atoms at positions -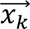 with masses *m*_k_, we can write two momentum conservation laws as

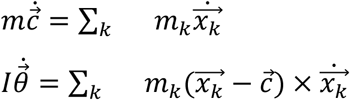

These conservation laws provide a linear relationship between *instantaneous* motions in the Cartesian and the rigid-body spaces. We can compactly represent them with linear projector operators written in a matrix form. Indeed, having a molecular system composed of *N* atoms, we can split it into *B* rigid blocks, which can be *B* residues in the case of a protein. Then, a 6*B* × 3*N* projector matrix *P* will provide a relationship between the two spaces. It is composed of *B* positional *P^c^* and orientational *P^θ^* sub-matrices of size 3 × 3*N_b_* each, where *N_b_* is the number of atoms in the *b*-th rigid body. Each 3 × 3 element of these matrices reads as

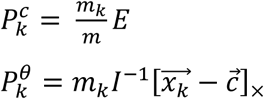

On the next step, we construct a 6*B* × 6*B* covariance matrix *C* in the RTB space from a normalized 3*N* × *t* matrix X collected from *t* MD frames,

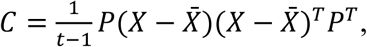

where matrix *X̅* corresponds to a mean molecular conformation over *t* frames. We decompose *C* as *C* = *VΛV^T^*, where *V* is a 6*B* × 6*B* matrix with each column defining an RTB eigenvector or a principal component that we interpret as an instantaneous linear and angular velocity of each rigid block (residue). *Λ* is a diagonal matrix containing the eigenvalues, which may depend on the sampling biases along the MD trajectory. Finally, we follow the original AFMFit strategy. Each eigenvector *V* has six components per residue: three correspond to the instantaneous linear velocities *v⃗* and the other three to the instantaneous angular velocities 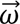. We extrapolate the coordinate *x⃗* of each atom in a residue at an amplitude *α* as

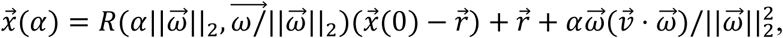

where we parameterize rigid rotation *R* with an angle-axis notation. Here, the center of rotation *r⃗* for a residue with the center of mass *c⃗* is given by

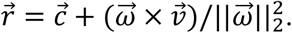

Following AFMFit, we also include into the calculations six zero-frequency eigenvectors corresponding to rigid-body translations and rotations. However, we compute them analytically. After this procedure, we follow the original flexible fitting step from AFMFit.

### MD Simulations

Molecular dynamics (MD) simulations were performed to generate libraries of conformations for Ca^2+^-bound and Ca^2+^-unbound alpha-actinin models. The initial models for alpha-actinin with and without Ca^2+^ in the bulk system were made from the PDB entry 1SJJ. The alpha-actinin was solvated in a 30 mM KCl solution. The system without Ca^2+^ consists of one alpha-actinin, 182 K^+^, 128 Cl^−^, and 251,756 water molecules. In the system with Ca^2+^, four K^+^ were removed and two Ca^2+^ were placed at the center of C_γ_ of Asp768 and Asp771. It was confirmed that Ca^2+^ ions are stably bound to Asp residues during the simulation. The histidine residues were assumed to be in the protonation state at pH = 7, since they face the bulk solution. The size of the system was 168.856 × 325.387 × 144.838 Å^3^. AMBER ff19SB^67^, TIP3P^68^, the Li and Merz 12-6^69^, the Li and Merz 12-6-4^70^ models were employed for the proteins, water molecules, monovalent ions, and divalent ions, respectively. The system was equilibrated by 1 ns MD simulations with constant temperature (300 K) and pressure conditions (1 bar), followed by 10 ns MD simulations with constant temperature (300 K) and volume conditions. All MD simulations were performed using AMBER22 (D. A. Case, H. M. Aktulga, K. Belfon, I. Y. Ben-Shalom, J. T. Berryman, S. R. Brozell, D. S. Cerutti, T. E. Cheatham, I. G. A. Cisneros, V. W. D. Cruzeiro, T. A. Darden, N. Forouzesh, G. Giambasų, T. Giese, M. K. Gilson, H. Gohlke, A. W. Goetz, J. Harris, S. Izadi, S. A. Izmailov, et al., AMBER22; University of California: San Francisco, 2022). A Berendsen thermostat^71^ was used to control the temperature, and a Monte Carlo barostat with anisotropic scaling was used to control the pressure. The SHAKE algorithm^72^ was used to keep the bond length having H atoms constant, enabling the use of a time step of 2 fs. The periodic boundary condition was imposed, and long-range interactions were calculated by the particle mesh Ewald method^73^ with a 10 Å cutoff in real space. After the equilibration, the MD simulation was performed for 1µs with constant temperature at 300 K and volume conditions using the Berendsen thermostat^71^. The trajectories were saved at each 1 ns to generate 1000 configuration libraries, which were used to analyse principal component (PC) modes using principal component analysis (PCA) to capture the bending and twisting angles, and distances between domains.

### Principal component analysis

Before the PCA, the configuration of alpha-actinin was aligned assuming that C_α_ atoms in the center part of the rod domain (residue 485-525) have the lowest root mean square fluctuation (RMSF). To compare the conformational library of AFMfit-NMA and MD simulations for Ca^2+^-bound and Ca^2+^-unbound states, we chose to reduce the dimensionality of the libraries using PCA. We normalized the number of elements in the six conformational libraries of AFMfit-NMA, AFMfit-MD, and MD simulations for Ca^2+^-bound and Ca^2+^-unbound states, ensuring each library contains 1,000 conformations. We combined these libraries to obtain a total library of 6,000 conformations. These conformations were then projected onto the same subspace using PCA. Then PCA was performed using singular value decomposition (SVD)^74^ on the mean-subtracted C_α_ coordinates of the combined libraries. We reduced the dimension to three principal components by retaining only the principal components (PCs) that have the most impact on the variance, more precisely the PCs that have an explained variance ratio (i.e. the normalized squared singular values) of more than 10%. To obtain a subspace on a per-atom basis, the eigenvectors were normalized by the number of C_α_ atoms in the system.

### Analyses of domain distances and twisting and bending angles in alpha-actinin

The ABD was divided into two domains: CH1, residue 26-138; CH2, residue 139-251. EF12 is residue 749-818, and EF34 is residue 819-888. The center of C_α_ of each domain was used to compute domain distances. Since there are four centers, there are six distances.

For the analyses of bending and twisting angles, the alpha-actinin was aligned as described in PCA. The C_α_ atoms in residue 485-525 were placed on the xy-plane and their average coordinate was set to be the origin. The rod domain was placed along the x-axis (**Fig. 6A**). The position of the center of C_α_ atoms of CH1 and CH2 in one chain and EF12 and EF34 in the other chain is denoted as (x_1_, y_1_, z_1_) (end 1), and that of the other end (end 2) is denoted as (x_2_, y_2_, z_2_). x_1_ is chosen so that x_1_ > 0, so x_2_ < 0. The bending angle for end 1 was calculated by tan^−1^(z_1_/x_1_), and that for end 2 by −tan^−1^(z_2_/x_2_). In this definition, the bending angles in alpha-actinin (PDB ID: 1SJJ) were −1.4 and 8.5 degrees.

For the twisting angle calculation, the position of the center of Cα atoms of CH1 and CH2 at end 1 is denoted as (x_1c_, y_1c_, z_1c_), and that of EF12 and EF34 at end 1 is denoted as (x_1e_, y_1e_, z_1e_). The twisting angle for end 1 was defined as the angle between a vector from the center of EF12 and EF34 to the center of CH1 and CH2 in the yz-plane and y axis (**Fig. 6B**): that is, tan^−1^((z_1c_ − z_1e_)/(y_1c_ − y_1e_)). For end 2, considering the orientation, it was calculated by tan^−1^(−(z_2c_ − z_2e_)/(y_2c_ − y_2e_)). When the twisting angle became less than −100 degrees, 360 degrees was added to the angle, since it is a part of the fluctuation over 180 degrees. In this definition, the twisting angles in alpha-actinin (PDB ID: 1SJJ) were −12.8 and 20.5 degrees.

## Data availability

All the data are included in the Supplementary Information or as source data. The conformation libraries, available in PDB format, can be accessed upon reasonable request.

## Code availability

https://gricad-gitlab.univ-grenoble-alpes.fr/GruLab/AFMfit-NMA

## Acknowledgments

The authors express their gratitude for the facility and financial support provided by the WPI Nano Life Science Institute (WPI NanoLSI), Kanazawa University; the instrumental assistance provided by Toshio Ando, Noriyuki Kodera, and Hiroki Konno (WPI NanoLSI). The authors also acknowledge the financial support from KAKENHI (Japan Society for the Promotion of Science) for K.X.N. (19K06581, 23K05713, 23H02452-01) and T.S. (24K01308). The MD simulations were carried out on the supercomputers at the Research Center for Computational Science in Okazaki, Japan (Project: 24-IMS-096). This study was also partially supported by the French National Research Agency (research grant ANR-23-CE45-0024-01) and within the framework of the “Investissements d’avenir” program (ANR-15-IDEX-02) for S.G.

## Author Contributions

K.X.N. designed the research; K.X.N., T.S., R.V. and S.G. performed the research; K.X.N., T.S., R.V., H.G.N., N.T.P.L., and S.G. analyzed the data; K.X.N., T.S., R.V. and S.G. wrote the paper.

---

**Supplementary Figure 1.**
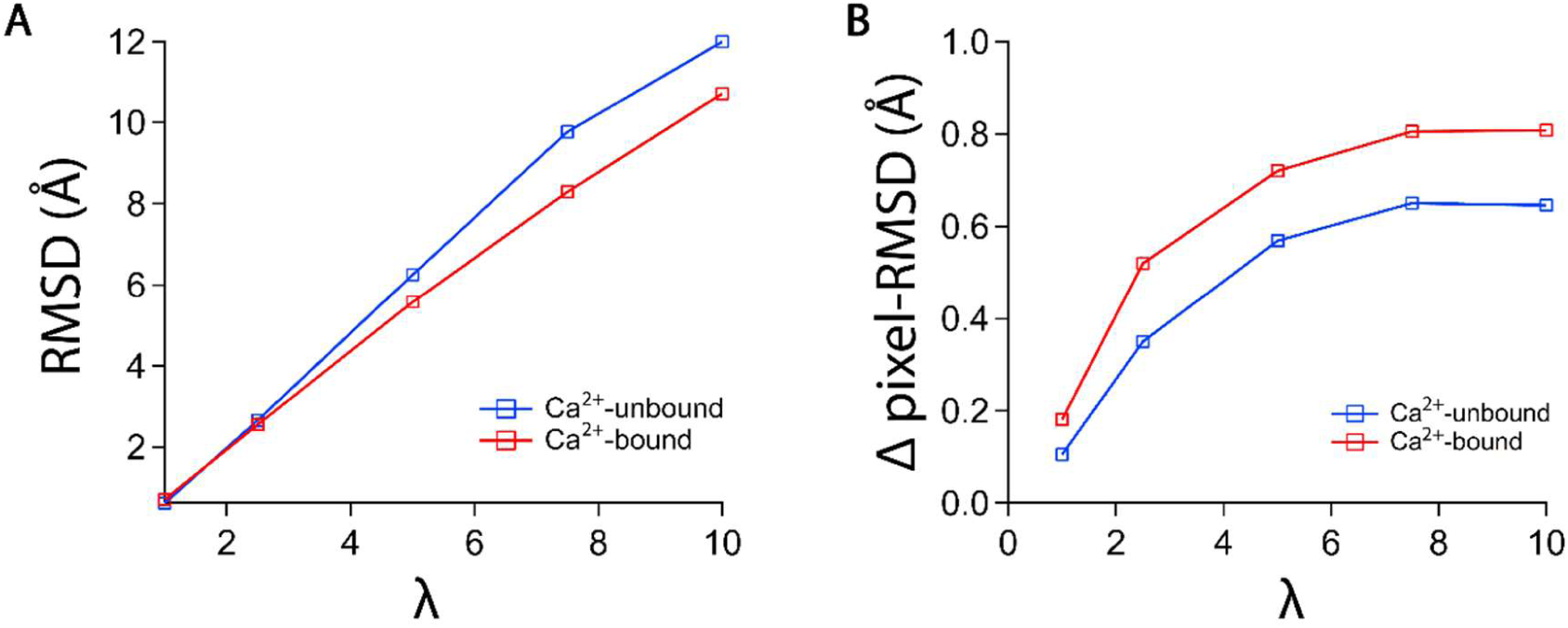
Optimization of the flexible fitting parameter (λ) The flexible fitting balanced structural preservation and data similarity, controlled by λ. An exhaustive search on sample particles identified the optimal λ = 7.5, yielding an average RMSD of ∼10 Å from the initial model, consistent with conformational motions observed in MD simulations of α-actinin.

**Supplementary Video 1** HS-AFM imaging of structure and dynamic conformational changes of Ca^2+^-bound and Ca^2+^-unbound alpha-actinin in solution. The AFM images were smoothed with a Gaussian filter (σ = 1). Imaging rate was done at 100 ms per frame. The Supplementary Video plays at 5 fps. Scale bars: 25 nm.

**Supplementary Video 2** The HS-AFM Supplementary Videos of Ca²⁺-bound alpha-actinin (left) and the atomic model (right) that were obtained through rigid and flexible fitting using AFMfit-NMA.

**Supplementary Video 3** The HS-AFM Supplementary Videos of Ca²⁺-unbound alpha-actinin (left) and the atomic model (right) that were obtained through rigid and flexible fitting using AFMfit-NMA.

**Supplementary Video 4** PC1 captured the ensemble of bending motions in the x-axis of alpha-actinin.

**Supplementary Video 5** PC2 captured the ensemble of bending motions in the y-axis of alpha-actinin.

**Supplementary Video 6** PC3 captured the ensemble of twisting motions of alpha-actinin.

**Supplementary Table 1.** Summary of the height and length (mean ± SD) distributions of ABDs in Ca²⁺-bound and Ca^2+^-unbound alpha-actinin analyzed using conformations obtained from HS-AFM. Statistical differences were assessed using a two-population *t-test*, with a significant difference confirmed at *p* ≤ 0.05. Related to **Figure 2**.

|  | Height (nm) | Length (nm) |
| --- | --- | --- |
|  | (N) | (N) |
| Ca<sup>2+</sup>-bound state | 5.7 ± 0.9<br>(6080) | 25.1 ± 2.3<br>(3040) |
| Ca<sup>2+</sup>-unbound state | 4.8 ± 0.7<br>(3885) | 25.0 ± 3.2<br>(1941) |
| *p*-value (*t*-test) | 0 | 0.78736 |

**Supplementary Table 2.** Summary of the distances between domains (mean ± SD) in Ca^2+^-bound and Ca^2+^-unbound alpha-actinin, derived from conformational libraries obtained from AFMfit-NMA, AFMfit-MD, and MD simulations. The domain distances were measured separately for chain 1 and chain 2 in individual conformations (stored as PDB files) for both Ca^2+^-bound and Ca^2+^-unbound states. The final data represent the averaged values of the two chains. A two-population *t-test* was conducted to evaluate the statistical difference (*p* ≤ 0.05) between the mean values of the Ca^2+^-bound and Ca^2+^-unbound states. **Related to** Figure 5.

| Domain distance<br>(nm) | Ca<sup>2+</sup>-bound/<br>AFMfit-NMA | Ca<sup>2+</sup>-unbound/<br>AFMfit-NMA | *t*-test (*p*-value) |
| --- | --- | --- | --- |
| CH1-CH2 | 2.45 ± 0.01 | 2.46 ± 0.01 | 4.6 × 10<sup>-10</sup> |
| CH1-EF12 | 3.89 ± 0.16 | 3.76 ± 0.21 | 1.9 × 10<sup>-62</sup> |
| CH1-EF34 | 3.06 ± 0.01 | 3.07 ± 0.02 | 8.7 × 10<sup>-38</sup> |
| CH2-EF12 | 4.58 ± 0.06 | 4.53 ± 0.09 | 1.8 × 10<sup>-49</sup> |
| CH2-EF34 | 3.63 ± 0.01 | 3.63 ± 0.02 | 1.5 × 10<sup>-08</sup> |
| EF12-EF34 | 2.81 ± 0.05 | 2.78 ± 0.07 | 7.7 × 10<sup>-48</sup> |
| N | 1466 | 1140 |  |
| Domain distance<br>(nm) | Ca<sup>2+</sup>-bound/<br>AFMfit-MD | Ca<sup>2+</sup>-unbound/<br>AFMfit-MD | *t*-test (*p*-value) |
| CH1-CH2 | 2.48 ± 0.07 | 2.51 ± 0.11 | 6.8x10<sup>-14</sup> |
| CH1-EF12 | 3.68 ± 0.23 | 3.80 ± 0.51 | 4.0x10<sup>-12</sup> |
| CH1-EF34 | 2.54 ± 0.13 | 3.23 ± 0.23 | 0 |
| CH2-EF12 | 4.20 ± 0.20 | 4.54 ± 0.26 | 2.1x10<sup>-191</sup> |
| CH2-EF34 | 3.35 ± 0.16 | 3.58 ± 0.20 | 2.9x10<sup>-152</sup> |
| EF12-EF34 | 2.81 ± 0.24 | 2.78 ± 0.21 | 2.7x10<sup>-3</sup> |
| N | 1000 | 1000 |  |
| Domain distance<br>(nm) | Ca<sup>2+</sup>-bound/<br>MD | Ca<sup>2+</sup>-unbound/<br>MD | *t*-test (*p*-value) |
| CH1-CH2 | 2.57 ± 0.02 | 2.51 ± 0.11 | 1.5x10<sup>-69</sup> |
| CH1-EF12 | 3.94 ± 0.19 | 3.80 ± 0.51 | 2.5x10<sup>-17</sup> |
| CH1-EF34 | $2.74 \pm 0.09$ | $3.23 \pm 0.23$ | 0 |
| CH2-EF12 | $4.49 \pm 0.07$ | $4.54 \pm 0.26$ | $5.3 \times 10^{-9}$ |
| CH2-EF34 | $3.59 \pm 0.08$ | $3.58 \pm 0.20$ | 0.06059 |
| EF12-EF34 | $3.14 \pm 0.12$ | $2.78 \pm 0.21$ | 0 |
| N | 1000 | 1000 |  |

**Supplementary Table 3.** Summary of the bending and twisting angles of the ABD (mean ± SD) near the neck domain relative to the rod domain in Ca^2+^-bound and Ca^2+^-unbound alpha-actinin, derived from conformational libraries obtained from AFMfit and MD simulations. A two-population *t-test* was conducted to evaluate the statistical difference (*p* ≤ 0.05) between the mean values of the Ca^2+^-bound and Ca^2+^-unbound states obtained from AFMfit or MD simulations. **Related to Figure 6**.

| **Figure 6A** | Bending angle<br>(degree) | N |
| --- | --- | --- |
| Chain 1-Ca<sup>2+</sup>-bound-AFMfit-NMA | 7.0 ± 6.7 | 1466 |
| Chain 2-Ca<sup>2+</sup>-bound-AFMfit-NMA | 4.3 ± 6.5 | 1466 |
| Chain 1-Ca<sup>2+</sup>-unbound-AFMfit-NMA | 1.7 ± 7.7 | 1140 |
| Chain 2-Ca<sup>2+</sup>-unbound-AFMfit-NMA | 2.8 ± 8.9 | 1140 |
| Chain 1-Ca<sup>2+</sup>-bound-AFMfit-MD | 7.0 ± 4.5 | 1000 |
| Chain 2-Ca<sup>2+</sup>-bound-AFMfit-MD | 3.6 ± 3.9 | 1000 |
| Chain 1-Ca<sup>2+</sup>-unbound-AFMfit-MD | 6.0 ± 5.3 | 1000 |
| Chain 2-Ca<sup>2+</sup>-unbound-AFMfit-MD | 1.0 ± 6.7 | 1000 |
| Chain 1-Ca<sup>2+</sup>-bound-MD | 8.0 ± 4.5 | 1000 |
| Chain 2-Ca<sup>2+</sup>-bound-MD | 5.3 ± 4.3 | 1000 |
| Chain 1-Ca<sup>2+</sup>-unbound-MD | 11.3 ± 5.5 | 1000 |
| Chain 2-Ca<sup>2+</sup>-unbound-MD | 4.7 ± 6.1 | 1000 |
| **Figure 6B** | Twisting angle<br>(degree) | N |
| Chain 1-Ca<sup>2+</sup>-bound-AFMfit-NMA | 16.7 ± 24.6 | 1466 |
| Chain 2-Ca<sup>2+</sup>-bound-AFMfit-NMA | 101.2 ± 37.5 | 1466 |
| Chain 1-Ca<sup>2+</sup>-unbound-AFMfit-NMA | 7.5 ± 29.4 | 1140 |
| Chain 2-Ca<sup>2+</sup>-unbound-AFMfit-NMA | 128.6 ± 41.5 | 1140 |
| Chain 1-Ca<sup>2+</sup>-bound-AFMfit-MD | 57.7 ± 14.5 | 1000 |
| Chain 2-Ca<sup>2+</sup>-bound-AFMfit-MD | 53.7 ± 19.8 | 1000 |
| Chain 1-Ca<sup>2+</sup>-unbound-AFMfit-MD | 6.3 ± 33.2 | 1000 |
| Chain 2-Ca<sup>2+</sup>-unbound-AFMfit-MD | 93.6 ± 32.1 | 1000 |
| Chain 1-Ca<sup>2+</sup>-bound-MD | 53.4 ± 16.2 | 1000 |
| Chain 2-Ca<sup>2+</sup>-bound-MD | 61.5 ± 12.5 | 1000 |
| Chain 1-Ca<sup>2+</sup>-unbound-MD | 28.5 ± 11.4 | 1000 |
| Chain 2-Ca<sup>2+</sup>-unbound-MD | 74.9 ± 15.3 | 1000 |

## References

1. Ando, T., Fukuda, S., Ngo, K. X. & Flechsig, H. High-Speed Atomic Force Microscopy for Filming Protein Molecules in Dynamic Action. Annu. Rev. Biophys. 53, 19–39 (2024).

2. Ando, T. et al. A high-speed atomic force microscope for studying biological macromolecules. Proc. Natl. Acad. Sci. U. S. A. 98, 12468–12472 (2001).

3. Shimizu, M. et al. An ultrafast piezoelectric Z-scanner with a resonance frequency above 1.1 MHz for high-speed atomic force microscopy. Rev. Sci. Instrum. 93, 013701 (2022).

4. Ando, T., Uchihashi, T. & Fukuma, T. High-speed atomic force microscopy for nano-visualization of dynamic biomolecular processes. Prog. Surf. Sci. 83, 337– 437 (2008).

5. Ando, T. et al. High-speed AFM and nano-visualization of biomolecular processes. Pflugers Arch. Eur. J. Physiol. 456, 211–225 (2008).

6. Ngo, K. X. et al. Allosteric regulation by cooperative conformational changes of actin filaments drives mutually exclusive binding with cofilin and myosin. Sci. Rep. 6, 1–11 (2016).

7. Ngo, K. X., Kodera, N., Katayama, E., Ando, T. & Uyeda, T. Q. Cofilin-induced unidirectional cooperative conformational changes in actin filaments revealed by high-speed atomic force microscopy. Elife 4, 1–22 (2015).

8. Ngo, Kien Xuan, Vu, H. T., Umeda, K., Trinh, M. N., Kodera, N. & Uyeda, T. Deciphering the actin structure-dependent preferential cooperative binding of cofilin. Elife 13, 1–37 (2024).

9. Yoshioka, Y. et al. PQBP3 prevents senescence by suppressing PSME3-mediated proteasomal Lamin B1 degradation. EMBO J. 43, 3968–3999 (2024).

10. Kodera, N. et al. Structural and dynamics analysis of intrinsically disordered proteins by high-speed atomic force microscopy. Nat. Nanotechnol. 16, 181–189 (2021).

11. Fukuda, S. & Ando, T. Faster high-speed atomic force microscopy for imaging of biomolecular processes. Rev. Sci. Instrum. 92, (2021).

12. Zwier, M. C. & Chong, L. T. Reaching biological timescales with all-atom molecular dynamics simulations. Curr. Opin. Pharmacol. 10, 745–752 (2010).

13. Hollingsworth, S. A. & Dror, R. O. Molecular dynamics simulation for all. Neuron 99, 1–29 (2018).

14. Karplus, M. & McCammon, J. A. Molecular dynamics simulations of biomolecules. Nat. Struct. Biol. 9, 646–652 (2002).

15. Kovermann, M., Rogne, P. & Wolf-Watz, M. Protein dynamics and function from solution state NMR spectroscopy. Q. Rev. Biophys. 49, e6 (2016).

16. Hu, Y. et al. NMR-Based Methods for Protein Analysis. Anal. Chem. 93, 1866– 1879 (2021).

17. Golji, J., Collins, R. & Mofrad, M. R. K. Molecular mechanics of the α-actinin rod domain: Bending, torsional, and extensional behavior. PLoS Comput. Biol. 5, (2009).

18. Adcock, S. A. & McCammon, J. A. Molecular dynamics: Survey of methods for simulating the activity of proteins. Chem. Rev. 106, 1589–1615 (2006).

19. Sumino, A., Sumikama, T., Uchihashi, T. & Oiki, S. High-speed AFM reveals accelerated binding of agitoxin-2 to a K+ channel by induced fit. Sci. Adv. 5, 1–10 (2019).

20. Chu, J. W. & Voth, G. A. Allostery of actin filaments: Molecular dynamics simulations and coarse-grained analysis. Proc. Natl. Acad. Sci. U. S. A. 102, 13111–13116 (2005).

21. Li, M. & Zheng, W. All-atom molecular dynamics simulations of actin-myosin interactions: A comparative study of cardiac α myosin, β myosin, and fast skeletal muscle myosin. Biochemistry 52, 8393–8405 (2013).

22. Yu, I. et al. Biomolecular interactions modulate macromolecular structure and dynamics in atomistic model of a bacterial cytoplasm. Elife 5, 1–22 (2016).

23. Lindorff-Larsen, K., Piana, S., Dror, R. O. & Shaw, D. E. How fast-folding proteins fold. Science (80-.). 334, 517–520 (2011).

24. Vakser, I. A., Grudinin, S., Jenkins, N. W., Kundrotas, P. J. & Deeds, E. J. Docking-based long timescale simulation of cell-size protein systems at atomic resolution. Proc. Natl. Acad. Sci. U. S. A. 119, 1–8 (2022).

25. Feig, M. & Sugita, Y. Whole-cell models and simulations in molecular detail. Annu. Rev. Cell Dev. Biol. 35, 191–211 (2019).

26. Hoffmann, A. & Grudinin, S. NOLB: Nonlinear Rigid Block Normal-Mode Analysis Method. J. Chem. Theory Comput. 13, 2123–2134 (2017).

27. Tama, F. & Sanejouand, Y. H. Conformational change of proteins arising from normal mode calculations. Protein Eng. 14, 1–6 (2001).

28. Tama, F., Valle, M., Frankt, J. & Brooks, C. L. Dynamic reorganization of the functionally active ribosome explored by normal mode analysis and cryo-electron microscopy. Proc. Natl. Acad. Sci. U. S. A. 100, 9319–9323 (2003).

29. Fukami, K. et al. Requirement of phosphatidylinositol 4,5-bisphosphate for α-actinin function. Nature 359, 150–152 (1992).

30. Fraley, T. S. et al. Phosphoinositide binding inhibits α-actinin bundling activity. J. Biol. Chem. 278, 24039–24045 (2003).

31. Taylor, K. a & Taylor, D. W. Formation of two-dimensional complexes of F-actin and crosslinking proteins on lipid monolayers: demonstration of unipolar alpha-actinin-F-actin crosslinking. Biophys. J. 67, 1976–83 (1994).

32. Young, P. The interaction of titin and alpha-actinin is controlled by a phospholipid-regulated intramolecular pseudoligand mechanism. EMBO J. 19, 6331–6340 (2000).

33. Ribeiro, E. D. A. et al. The structure and regulation of human muscle α-Actinin. Cell 159, 1447–1460 (2014).

34. Liu, J., Taylor, D. W. & Taylor, K. A. A 3-D reconstruction of smooth muscle α-actinin by CryoEm reveals two different conformations at the actin-binding region. J. Mol. Biol. 338, 115–125 (2004).

35. Foley, K. S. & Young, P. W. The non-muscle functions of actinins: An update. Biochem. J. 459, 1–13 (2014).

36. Pinotsis, N. et al. Calcium modulates the domain flexibility and function of an α-actinin similar to the ancestral a-actinin. Proc. Natl. Acad. Sci. U. S. A. 117, 22101–22112 (2020).

37. Tang, J., Taylor, D. W. & Taylor, K. a. The three-dimensional structure of alpha-actinin obtained by cryoelectron microscopy suggests a model for Ca(2+)-dependent actin binding. J. Mol. Biol. 310, 845–58 (2001).

38. Vuillemot, R., Pellequer, J. L. & Grudinin, S. Deciphering conformational dynamics in AFM data using fast nonlinear NMA and FFT-based search with AFMFit. *Commun*. Biol. 8, 1–13 (2025).

39. Shams, H., Golji, J., Garakani, K. & Mofrad, M. R. K. Dynamic Regulation of α-Actinin’s Calponin Homology Domains on F-Actin. Biophys. J. 110, 1444–1455 (2016).

40. Uchihashi, T., Kodera, N. & Ando, T. Guide to video recording of structure dynamics and dynamic processes of proteins by high-speed atomic force microscopy. Nat. Protoc. 7, 1193–1206 (2012).

41. Kodera, N., Yamamoto, D., Ishikawa, R. & Ando, T. Video imaging of walking myosin V by high-speed atomic force microscopy. Nature 468, 72–76 (2010).

42. Voss, J. M., Harder, O. F., Olshin, P. K., Drabbels, M. & Lorenz, U. J. Rapid melting and revitrification as an approach to microsecond time-resolved cryo-electron microscopy. Chem. Phys. Lett. 778, 138812 (2021).

43. Bongiovanni, G., Harder, O. F., Drabbels, M. & Lorenz, U. J. Microsecond melting and revitrification of cryo samples with a correlative light-electron microscopy approach. Front. Mol. Biosci. 9, 1–7 (2022).

44. Lorenz, U. J. Microsecond time-resolved cryo-electron microscopy. Curr. Opin. Struct. Biol. 87, 102840 (2024).

45. Vuillemot, R., Miyashita, O., Tama, F., Rouiller, I. & Jonic, S. NMMD: Efficient Cryo-EM Flexible Fitting Based on Simultaneous Normal Mode and Molecular Dynamics atomic displacements. J. Mol. Biol. 434, 167483 (2022).

46. Boehr, D. D., Dyson, H. J. & Wright, P. E. An NMR perspective on enzyme dynamics. Chem. Rev. 106, 3055–3079 (2006).

47. Wolf-Watz, M. et al. Linkage between dynamics and catalysis in a thermophilic-mesophilic enzyme pair. Nat. Struct. Mol. Biol. 11, 945–949 (2004).

48. Purslow, J. A., Khatiwada, B., Bayro, M. J. & Venditti, V. NMR Methods for Structural Characterization of Protein-Protein Complexes. Front. Mol. Biosci. 7, 1–8 (2020).

49. Camacho-Zarco, A. R. et al. NMR Provides Unique Insight into the Functional Dynamics and Interactions of Intrinsically Disordered Proteins. Chem. Rev. 122, 9331–9356 (2022).

50. Moraes, A. H. & Valente, A. P. Conformational dynamics and kinetics of protein interactions by nuclear magnetic resonance. J. Magn. Reson. Open 14–15, 100093 (2023).

51. Murata, K. & Wolf, M. Cryo-electron microscopy for structural analysis of dynamic biological macromolecules. Biochim. Biophys. Acta - Gen. Subj. 1862, 324–334 (2018).

52. Gauto, D. F. et al. Integrated NMR and cryo-EM atomic-resolution structure determination of a half-megadalton enzyme complex. Nat. Commun. 10, 1–12 (2019).

53. Schanda, P. & Haran, G. NMR and Single-Molecule FRET Insights into Fast Protein Motions and Their Relation to Function. Annu. Rev. Biophys. 53, 247–273 (2024).

54. Lorenz, M., Popp, D. & Holmes, K. C. Refinement of the F-actin model against x-ray fiber diffraction data by the use of a directed mutation algorithm. J. Mol. Biol. 234, 826–836 (1993).

55. Oda, T., Iwasa, M., Aihara, T., Maéda, Y. & Narita, A. The nature of the globular-to fibrous-actin transition. Nature 457, 441–445 (2009).

56. Fujii, T., Iwane, A. H., Yanagida, T. & Namba, K. Direct visualization of secondary structures of F-actin by electron cryomicroscopy. Nature 467, 724–728 (2010).

57. Murakami, K. et al. Structural basis for actin assembly, activation of ATP hydrolysis, and delayed phosphate release. Cell 143, 275–287 (2010).

58. González, M. A. Force fields and molecular dynamics simulations. Collect. SFN 12, 169–200 (2011).

59. Chmiela, S., Sauceda, H. E., Müller, K. R. & Tkatchenko, A. Towards exact molecular dynamics simulations with machine-learned force fields. Nat. Commun. 9, 1–10 (2018).

60. Bhavna, R. & Sonawane, M. A deep learning framework for quantitative analysis of actin microridges. *npj Syst*. Biol. Appl. 9, 1–15 (2023).

61. Degenhardt, M. F. S. et al. Determining structures of RNA conformers using AFM and deep neural networks. Nature 637, (2025).

62. Vinh, T., Nguyen, T., Ly, N. Q., Thi, N. & Le, P. AFMnanoSALQ: An Accurate Detection Framework for Semi-Automatic Labeling and Quantitative Analysis of α-Hemolysin Nanopores Using Intensity-Height Cues in HS-AFM Data. bioRxiv (2025). doi:10.1101/2025.02.26.640237

63. Jiang, Y., Wang, Z. & Scheuring, S. A structural biology compatible file format for atomic force microscopy. Nat. Commun. 16, 1–16 (2025).

64. Sumikama, T. et al. Computed Three-Dimensional Atomic Force Microscopy Images of Biopolymers Using the Jarzynski Equality. J. Phys. Chem. Lett. 13, 5365–5371 (2022).

65. Ngo, K. X. et al. Allosteric regulation by cooperative conformational changes of actin filaments drives mutually exclusive binding with cofilin and myosin. Sci. Rep. 6, 1–11 (2016).

66. Pu Liu, Dimitris K. Agrafiotis, D. L. T. Rapid Communication Fast Determination of the Optimal Rotational Matrix for Macromolecular Superpositions. J. Comput. Chem. 31, 1561–1563 (2010).

67. Tian, C. et al. Ff19SB: Amino-Acid-Specific Protein Backbone Parameters Trained against Quantum Mechanics Energy Surfaces in Solution. J. Chem. Theory Comput. 16, 528–552 (2020).

68. William L. Jorgensen, Jayaraman Chandrasekhar, and J. D. M. Comparison of simple potential functions for simulating liquid water. J. Chem. Phys. 79, 926–935 (1983).

69. Li, P., Song, L. F. & Merz, K. M. Systematic parameterization of monovalent ions employing the nonbonded model. J. Chem. Theory Comput. 11, 1645–1657 (2015).

70. Li, P. & Merz, K. M. Taking into account the ion-induced dipole interaction in the nonbonded model of ions. J. Chem. Theory Comput. 10, 289–297 (2014).

71. Berendsen, H. J. C., Postma, J. P. M., Van Gunsteren, W. F., Dinola, A. & Haak, J. R. Molecular dynamics with coupling to an external bath. J. Chem. Phys. 81, 3684–3690 (1984).

72. Ryckaert, J. P., Ciccotti, G. & Berendsen, H. J. C. Numerical integration of the cartesian equations of motion of a system with constraints: molecular dynamics of n-alkanes. J. Comput. Phys. 23, 327–341 (1977).

73. Essmann, U. et al. A smooth particle mesh Ewald method. J. Chem. Phys. 103, 8577–8593 (1995).

74. Wall, M. E., Rechtsteiner, A. & Rocha, L. M. Singular Value Decomposition and Principal Component Analysis. in A Practical Approach to Microarray Data Analysis 91–109 (2005). doi:10.1007/0-306-47815-3_5

75. Amyot, R. & Flechsig, H. BioAFMviewer: An interactive interface for simulated AFM scanning of biomolecular structures and dynamics. PLoS Comput. Biol. 16, 1–12 (2020).

